# Occurrence of Carbapenem-resistant *Enterobacterales* in swine wastewater in Shandong Province, China

**DOI:** 10.64898/2025.12.02.691800

**Authors:** Ying Chu, Yujie Miao, Liya Huang, Rong Wang, Yuxin Wang, Fengting Liao, Xiang Luo, Shuancheng Bai, Yubao Li

**Author notes:** Corresponding author at: College of Smart Agriculture, Yulin Normal University, Yulin, 53700, China. E-mail address (S.C. Bai); Liaocheng University, Liaocheng, Shandong 252000, China. E-mail address (Y. B. Li).

## Abstract

Swine wastewater can accelerate the spread of antibiotic-resistant bacteria among animals, the environment, and humans. Therefore, the molecular epidemiological characteristics of carbapenemase-producing *E. coli* isolates from swine wastewater was investigated. In this study, a total of 100 no-cloning carbapenemase-producing (*bla*_NDM-1_, *bla*_NDM-5_ and *bla*_OXA-48_-like) *E. coli* isolates were recovered from 316 swine wastewater samples collected from 29 swine farm in Shandong, China. Of which, two isolates that co-harbored *bla*_OXA-48_-like and *mcr-1*, as well as five isolates that co-harbored *bla*_NDM-5_ and *tet(X4)*. All isolates were multidrug-resistant (MDR), with a majority exhibiting resistance to meropenem, ciprofloxacin, imipenem and gentamicin. Whole genome sequencing (WGS) analysis indicated that these isolates were belonged to 12 distinct sequence types (STs), with the most prevalent STs being ST10 and ST5299. Additionally, Phylogenomic analysis revealed that clonal spread of *bla*_NDM_-positive ST10 *E. coli* isolates at swine wastewater in multiple cities, including Binzhou, Dezhou, Heze and Weihai. Meanwhile, a notable divergence in SNPs was observed in isolates from this study and public database, which indicates that these *bla*_NDM_-positive *E. coli* isolates have high genetic diversity. In addition, WGS analysis further revealed that *bla*_NDM_ co-existed with other antibiotic resistance genes, conferring resistance to twelve classes of antimicrobials. This study underscores the importance of surveillance for *bla*_NDM_-harboring microbes in swine wastewater, and continuous monitoring in swine wastewater is essential for ensure human and environmental health.

**IMPORTANCE:** The study described the NDM-positive *E. coli* strains isolated from swine wastewater, where it was associated with antibiotic exposure. Notably, swine wastewater was increasingly considered to be related to CRE hosts. Pathogens originating from this source can be transmitted to humans via direct and indirect contact, as well as through the ingestion or inhalation of contaminated water or aerosols. In addition, under the selective pressure of antibiotics, the co-harbored of *bla*_OXA-48_-like and *mcr-1*, *bla*_NDM-5_ and *tet(X4)* genes within the same host bacterium signifies the emergence of an extremely MDR pathogen, posing a grave threat to public health. Consequently, the monitoring of *bla*_NDM_ in swine wastewater is critical for containing the transmission and persistence of carbapenem-resistant pathogens.

---

The presence of carbapenem-producing *Enterobacteriaceae* (CPEs) in swine wastewater is a cause of increasing concern due to its role in environmental contamination and potential dissemination into human healthcare settings. These organisms often carry genes encoding intra- and inter-cellular gene mobility, virulence, metal tolerance and antibiotics genes (1). Such a gene pool can potentially interact with and modify the gene pool of environmental bacteria, and vice versa (2), and microorganisms resulting from these interactions might pose a health threat to livestock and poultry and, ultimately, back to humans. Accordingly, CRE in swine wastewater need more attention.

Swine wastewater, as a major recipient of substantial amounts of antimicrobial residues, wastewater is recognized as both a primary source and transmission pathway for resistance determinants (3). In reality, antimicrobial agents used by humans are not fully degraded, and their metabolites are excreted into the environment, generating adverse impacts on wastewater contamination and public health. However, the overuse of β-lactam antibiotics has promoted the enrichment of CPEs in livestock and poultry (4). The most common mechanism of CPEs is the expression of β-lactam-hydrolysing enzymes e.g, such as New NDM and KPC, encoded by *bla*_NDM_ and *bla*_KPC_, respectively. Unlike *bla*_KPC_ mainly found in bacteria from human infections; *bla*_NDM_ can be found across both humans and animal sectors, the same as the swine wastewater environment (5–7). *bla*_NDM_ are encoded by genes located mostly on mobile genetic elements (MGEs) such as plasmids, integrons, and transposons (8). In addition to *bla*_NDM_, these MGEs often carry resistance genes for aminoglycosides, CTX-M, and quinolones extended spectrum beta-lactamase (ESBL) genes (9–11). After that, the antibiotic resistance genes can also persist in aquatic environments and this promotes further dissemination of ARGs between aquatic environment and aquatic animals.

Studies indicate that CPE transmission is correlated throughout the entire food animal production chain. The CPEs increases gradually along the chicken and pig breeding (4.70%/2.0%)–slaughtering (7.60%/22.40%)–retail (65.56%/34.26%) chains (12). The clonal spread of CPEs was found between farmed ducks and slaughtered duck meats even from different farms (13). Such strains were detectable in samples from farmland (10.3%, 8/78), vegetable fields (7.3%, 3/41), and environment of chicken farms (25.6%, 41/160) which had been left vacant for a long period of time, and these strains can persist (14). In addition, the recent discoveries of plasmid-mediated *bla*_NDM_ or *bla*_OXA-48_-like genes co-harboring *mcr* and/or the *tet(X)* among *Enterobacteriaceae*, predict a return to the pre-antibiotic era and pose a severe threat to public health, which requires urgent monitoring in terms of its prevalence.

In this context, the present study aimed to conduct the surveillance of MDR *E. coli* from swine wastewater in Shandong, China through the phenotypic and molecular evaluation of the co-harboring *bla*_NDM-5_ and *tet(X4)*, *bla*_OXA-48_-like and *mcr-1* gene cluster genes involved to assess the possible role of the swine wastewater as a reservoir and dissemination pathway of such resistance mechanisms.

## RESULTS

### Prevalence of carbapenemase-producing *E. coli* isolates

In this study, a total of 100 CPEs were recovered from 316 wastewater samples from swine wastewater in Shandong, China. A total of two types of CPE were identified in this study, including 98 *bla*_NDM_-positive strains and 2 *bla*_OXA-48_-like-positive strains. and no other carbapenemase-encoding genes were detected among these carbapenem-resistant isolates. Among the 2 *bla*_NDM_ variants, *bla*_NDM-5_ was the dominant, constituting (94/98, 95.92%), followed by *bla*_NDM-1_ (4/98, 4.08%). The highest detection rate of *bla*_NDM_-positive *E. coli* isolates was observed in Dongying (20/20, 100%), followed by Heze (28/36, 77.8%), Weifang (5/30, 57.1%) and Dezhou (19/71, 26.8%). In contrast, Taian (3/14, 21.4%), Yantai (7/29, 21.4%), Binzhou (2/10, 20%), Weihai (5/30, 16.7%), Jinan (3/28, 10.7%) and Liaocheng (5/64, 7.8%) displayed the lowest detection rates (Figure 1).

**Figure 1.**
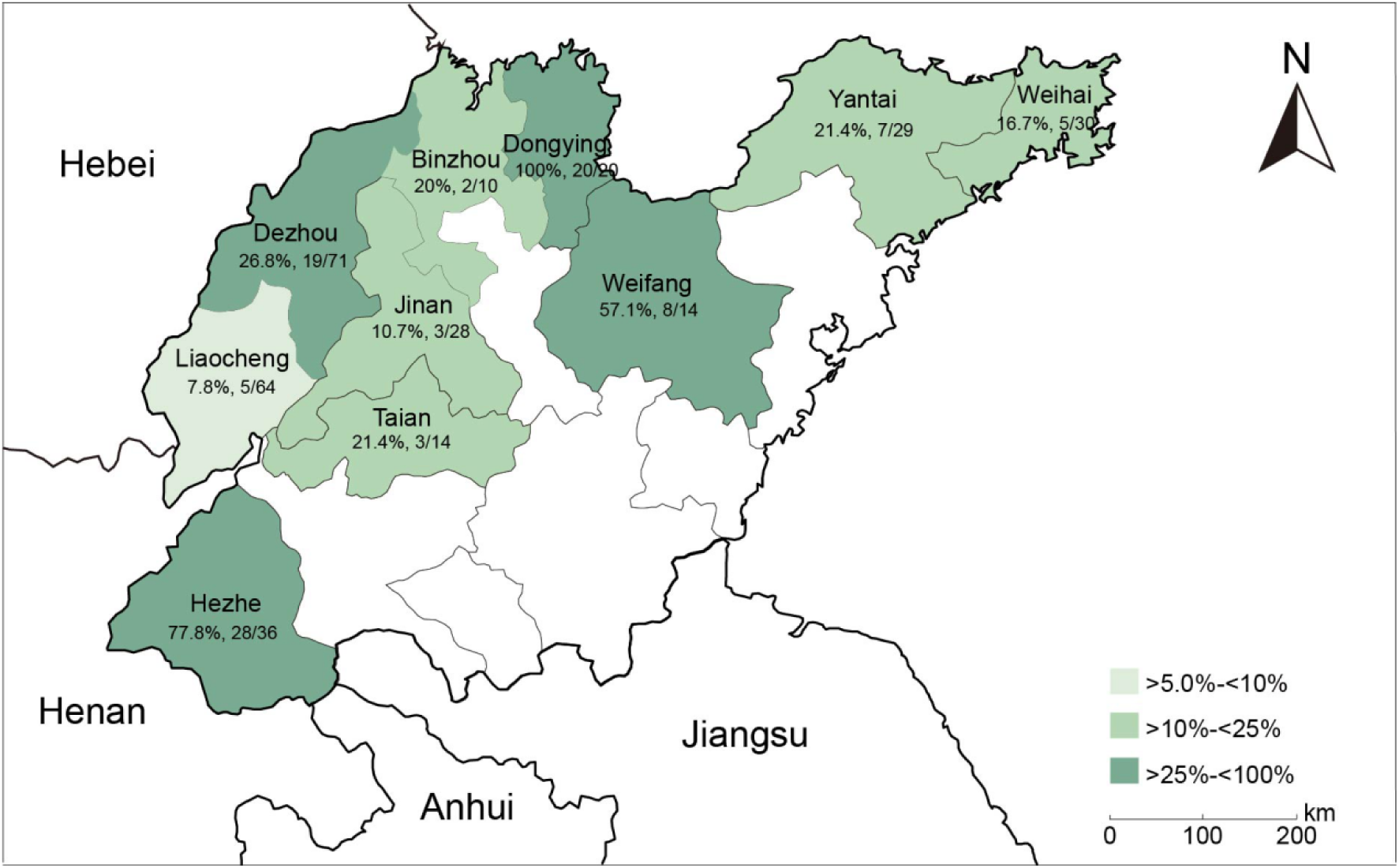
Geographical distribution of CPEs in Shandong Province, China.

### Antibiotic resistance phenotypes

We obtained all of 100 non-duplicate carbapenemase-producing *E. coli* strains exhibited resistance to meropenem, cefotaxime, ceftazidime, tetracycline, and trimethoprim-sulfamethoxazole (Figure 2, table S1). In addition, the majority of these isolates remained resistant to ciprofloxacin (76/100, 76%), imipenem (72/100, 72%) and gentamicin (45/100, 45%). In contrast, lower prevalence of resistance phenotypes was observed for colistin (10/100, 10%), amikacin (7/100, 7%), tigecycline (3/100, 3%), of note, none of isolates showed resistance to amtronam and fosfomycin.

**Figure 2.**
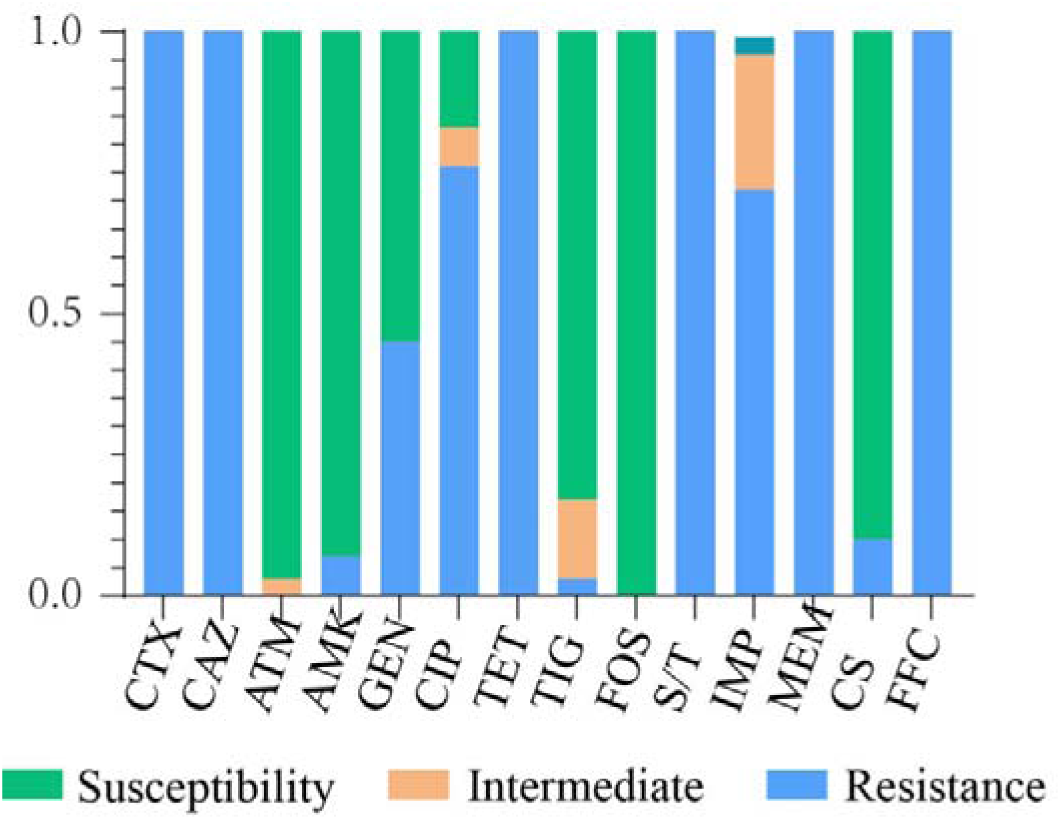
Minimum inhibitory concentrations of tested antimicrobial agents for the studied bacterial isolates. Note: CTX, cefotaxime; CAZ, ceftazidime; ATM, aztreonam; AMK, amikacin; GEN, gentamicin; CIP, ciprofloxacin; TET, tetracycline; TIG, tigecycline; FOS, fosfomycin; S/T, sulfamethoxazole/trimethoprim; IMP, imipenem; MEM, meropenem; CS, colistin; FFC, florfenicol.

### Phylogenetic analysis of NDM and OXA-48-like-positive *E. coli* isolates

Based on preliminary ERIC PCR analysis, 29 *E. coli* strains with distinct banding patterns were selected for whole-genome sequencing (WGS), and results revealed that these isolates were categorized into 12 distinct STs, with 4 isolates classified as unclassified STs. Overall, ST10 (20.7%, 6/29) was the most prevalent in Binzhou, Dezhou, Heze and Weihai, followed by ST5299 (17.2%, 5/29) in Dongying and Dezhou. These findings suggest a distinct predilection for specific geographic distributions (Figure 3, table S2). A phylogenetic tree was established using these CPEs, and revealed that all the *E. coli* isolates were classified into 3 distinct lineages. In the lineages I mainly belonged to ST5409 and were sourced from Taian, with these isolates sharing only 0∼2 core-genome SNP (cgSNP) among themselves. It is worth noting that all isolates in the lineages II mainly belonged to ST5229 and were sourced from Dongying and Dezhou. However, they exhibited significant SNP differences (1314 SNP), and the same as the strains from Dezhou carried *bla*_OXA-48_-like gene sharing 36 cgSNP. In addition, isolates in lineages III mainly belonged to ST10 and originated from Weifang, Bingzhou, Dezhou and Heze, and 3 new strains that failed to be classified as ST sharing only one SNP from the strains in Tai’an and Weifang. Our phylogenomic analysis demonstrated that the majority of *bla*_NDM_-positive *E. coli* isolates exhibited a significant degree of variation in core-genome sequences, suggesting a high genetic diversity of *bla*_NDM_-positive *E. coli* isolates from swine wastewater in Shandong Province, China.

**Figure 3.**
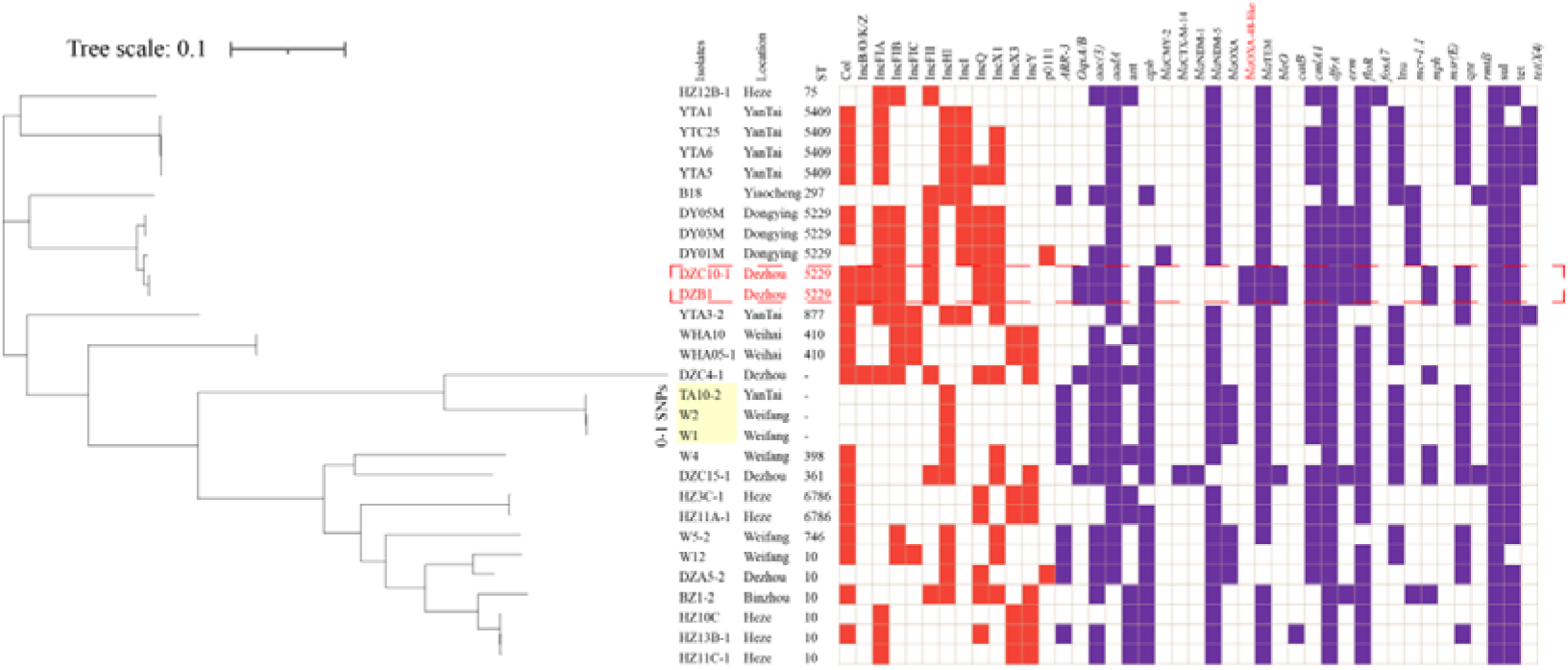
Phylogenetic analysis of Carbapenemase-producing *E. coli* isolates in this study. Bayesian evolutionary tree was constructed using core-genome SNPs. Each isolate is labeled with the city of isolation year and ST. The red-filled squares indicate the possession of the indicated ARGs.

To further assess the relationship between the isolates from the current study and other resources in China. A total of 397 *bla*_NDM_-positive *E. coli* isolates, obtained from 32 provincial-level administrative divisions across China, were randomly collected via the National Center for Biotechnology Information (NCBI) database (as April 2024) (Figure 4, table S3). Then, a maximum likelihood phylogenetic tree was constructed using these 426 *bla*_NDM_-positive *E. coli* isolates, which were grouped into 6 clades, among which the 29 isolates from this study distributed across 5 clades. In addition, a notable divergence in SNPs was observed in *bla*_NDM_-positive *E. coli* isolates between those analyzed in this study and those sourced from the NCBI database, which indicates that these *bla*_NDM_-positive *E. coli* isolates have high genetic diversity.

**Figure 4.**
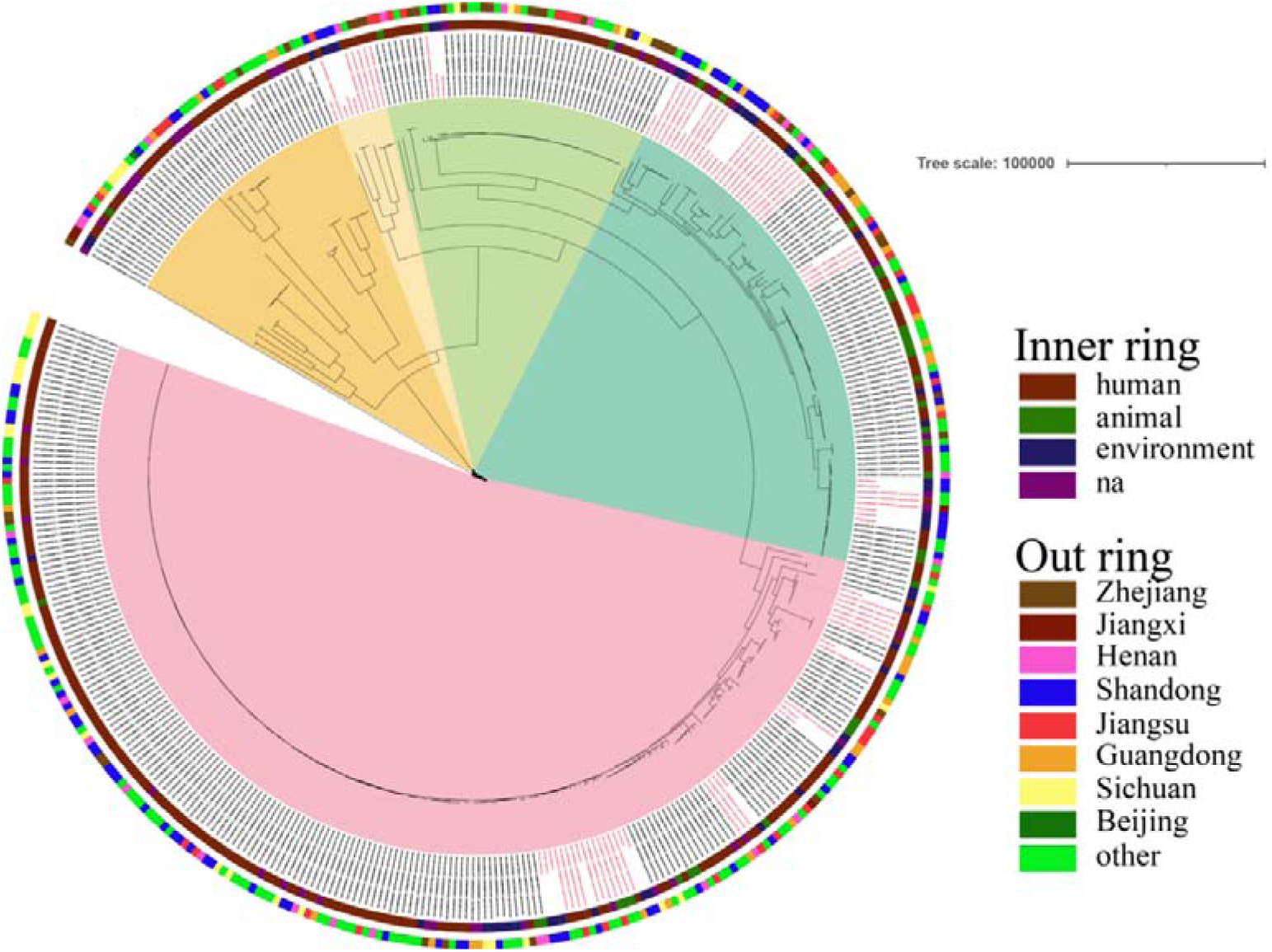
Phylogenetic structures of the*bla*_NDM-_ and *bla*_OXA-48_-like-positive *E. coli* isolates from this study and the GenBank database. Strain hosts and country of origin are indicated in the inner and outer rings, respectively.

### The genetic environments of *bla*_NDM_ and *bla*_OXA-48_-like

A total of 5 genetic contexts (type I to type V) were found in 27 *bla*_NDM_-positive isolates (Figure 5). The type IV genetic contexts was novel in the GenBank database. Notably, *bla*_NDM-5_ was found in 2 out of the 3 genetic context types. In all types, *bla*_NDM-5_ was directly associated with the *ble*_MBL_-*trpF*-*tat* genes. In addition, we found that the type I genomic context was the most prevalent *bla*_NDM_ genetic environment in this study (63%, 17/27), which is identical to a *bla*_NDM-5_-carrying plasmid of carbapenem-resistant *Enterobacteriaceae* isolates from humans in China (15). Meanwhile, the backbone structure of *bla*_NDM-5_ including the IS*Aba125*-IS*5*-*bla*_NDM-5_-*ble*_MBL_-*trpF*-*tat*-IS*6* was usually carrying the IncX3-type plasmid and highly conserved (16). In contrast, *bla*_NDM-1_ was linked to IS*5* and IS*3000* (Type IV and V). In type IV, *bla*_NDM-1_ is flanked by IS*3000* and IS*5* within the backbone region of IncY plasmids. We verified this result by PCR, using primers designed in the backbone region of the IncY plasmid and near *bla*_NDM-1_. In type V, the backbone carrying *bla*_NDM-1_ simultaneously harbors multiple virulence genes (*virB1-11* and virD4). Type II and Type III share a core resistance gene cluster (*ant(3’’)-Ia*-*qacE*-*sul*-*bla*_NDM-5_). However, Type III acquires unique evolutionary traits due to the insertion of the *yheS* gene (ABC transporter, ATP-binding protein) mediated by the Tn*3* transposon. In addition, two OXA-48-like-positive bacterial strains identified in this study share an identical genetic environment (*bla*_OXA-48_-like-IS*Kpn19*).

**Figure 5.**
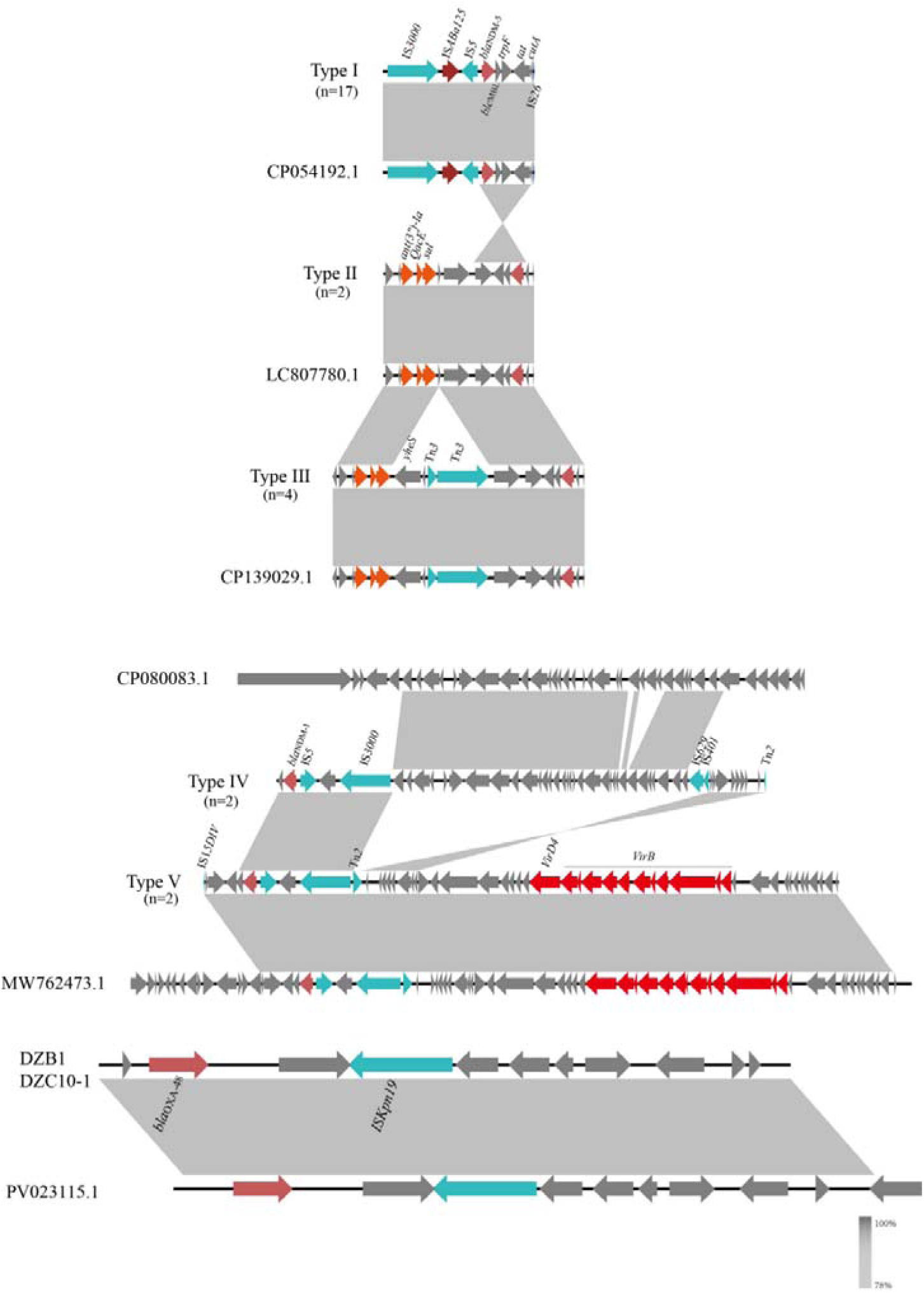
Genomic environments of *bla*_NDM_ and *bla*_OAX-48_-like of *E. coli* isolates. The figure was generated using Easyfig. Regions of homology are marked by shading and regions of ≥ 99.0 % nucleotide sequence identity are shaded grey. Arrows indicate the direction of transcription of the genes. In addition, red represents resistance gene, blue represents mobile genetic elements, black represents transfer gene tra clusters and yellow represents other coding sequence (CDS).

### Analysis of antibiotic resistance genes

We conducted a comprehensive antimicrobial analysis on all of the *E. coli* isolates that revealed the presence of the β-lactam resistance genes (*bla*_NDM-1_, *bla*_NDM-5_, *bla*_OXA-10_, *bla*_OXA-48_-like, *bla*_CTX-M_, *ble*_O_, *bla*_CMY-2_ and *bla*_TEM,_). Other important resistance determinants that conferring resistance to quinolones (*oqxA/B*, *qnr*), aminoglycosides (*rmtB*, *aadA, aph*, *ant* and *aac*), fosfomycin (*fosA*), chloramphenicol/florfenicol (*floR* and *catB*), sulfonamides (*sul*), macrolide (*erm*, *msr(E)* and *mph(A)*), rifampicin (*ARR-3*), phenicols (*cmlA*), lincomycin (*lnu*), tetracycline (*tet*) and trimethoprim (*dfrA*). Additionally, we identified two isolates that co-harbored *bla*_OXA-48_-like and *mcr-1*, as well as five isolates that co-harbored *bla*_NDM-5_ and *tet(X4)* (Figure 3).

### Plasmid analysis

A total of 13 incompatible group plasmid replicon types were detected among the 29 carbapenemase-producing *E. coli* isolates including Col (69%, 20/29), IncB/O/K/Z (10%, 3/29), IncFIA (52%, 15/29), IncFIB (41%, 12/29), IncFIC (14%, 4/29), IncFII (34%, 10/29), IncHI (52%, 15/29), IncI (31%, 9/29), IncQ (41%, 12/29), IncXI (52%, 15/29), IncX3 (24%, 7/29), IncY (34%, 10/29) and p0111 (7%, 2/29). It is worth noting that the cloning strains W1, W2 and TA10-2 from Weifang and Yantai only carry the IncHI plasmid, suggesting that this plasmid type represents a multidrug-resistant (MDR) plasmid harboring resistance determinants including *bla*_NDM_.

## DISCUSSION

In the present study, we investigated the prevalence of carbapenemases-producing *E. coli* isolates from swine wastewater in Shandong, China. Our results revealed a wide contamination of *bla*_NDM_- and *bla*_OXA-48_-like-producing *Enterobacteriaceae* from the environmental. Previous studies have also shown the detection of these species causing infections in the livestock production environment (17,18). Researches show that gram-negative bacteria, including *Enterobacteriaceae*, are able to survive on abiotic surfaces which present a pathogenic transmitter from one surface to another, or even from the environmental area to patients and to staff (19). and most likely exacerbate the transmission of *bla*_NDM_ and *bla*_OXA-48_-like.

More importantly, we detected *bla*_NDM_-positive *E. coli* with relatively high prevalence from environmental samples (31%, 98/316), and 2 OXA-48-like-producing *E. coli*. A high prevalence of *bla*_NDM_-positive and *bla*_OXA-48_-like positive was reported for all food animals with prevalence rates of chicken, cattle, and swine (3,20). Together with the current findings, these data strongly indicated that the Chinese poultry farm environment is likely to be an important reservoir for *bla*_NDM_ and *bla*_OXA-48_-like carrying bacteria. Importantly, carbapenems are not used in food animal production. Consequently, we surmise that the high prevalence of *bla*_NDM_-carrying bacteria in the present study may have originated from the use of amoxicillin on the farm, which probably provided a direct selection pressure for *bla*_NDM_ maintenance. This hypothesis has been supported by recent research evidence (4). In addition, the swine wastewater was an open space suitable, and this made it difficult to perform complete disinfection and sterilization of the farm environment. There was widespread contamination by *bla*_NDM_-positive *Enterobacteriaceae* in farm environments. In this study, *bla*_NDM-5_ was the predominant variant, consistent with previous reports identifying *E. coli* as the primary *Enterobacteriaceae* species carrying *bla*_NDM-5_ in both farm animals and their environments (6,21).

The genetic background analysis shows In type VI, was found a fusion plasmid was found that was recombined from an IncX3 plasmid carrying *bla*_NDM-5_ and an IncF plasmid carrying *ant(3’’)-Ia*, *qacE* and *sul*.This finding is consistent with our recent study, with the difference being that the previous study sampled retail chicken markets (22). It’s worth noting that *bla*_NDM-1_ is flanked by IS3000 and IS5 within the backbone region of IncY and IncX3 plasmids. A few reports of *bla*_NDM-1_ transmission via IncY plasmids, and there was only one report of *bla*_NDM-1_ transmission via IncY plasmids (23).

WGS analysis revealed that *bla*_NDM_ coexisted with other 29 other types of ARGs, 12 of which ARGs were highly prevalent with detection rates >50%. This diversity of ARGs confirms their role as potential sources of determinants of drug resistance (24). These ARGs encoded resistance to aminoglycosides, β-Lactam and tetracycline and can further exacerbate the spread of carbapenemases-resistant isolates in swine wastewater as well as among surrounding animals, humans and environments in pig farm. To date, indeed, there are only very limited findings on OXA-48 and OXA-48-like producing and *mcr-1* producing *E. coli* in livestock (25,26). In this study, WGS analysis revealed that *bla*_OXA-48_-like and *mcr-1* coexisted with other 14 other types of ARGs. The widespread presence of carbapenem-resistant *Enterobacteriaceae* (CRE) and *mcr*-positive *E. coli* poses a huge threat to both animal and human health (27). In fact, for now, the *tet(X4)*-positive isolates were mostly reported in animals such as pigs, chickens, cows and ducks, and *tet(X4)* has been previously identified in pigs and chickens at slaughters, from soil and dust in animal farms, and even pork from markets (28–30). However, despite the *tet(X4)* and *bla*_NDM-5_ genes being reported as co-harboured in *E coli* (31), it is unknown whether *tet(X4)* and *bla*_NDM-5_ could co-exist in ST5409 and ST877. Here we report the first identification of a carbapenem- and tigecycline-resistant *E. coli* isolate recovered from pig farm in China harbouring the *tet(X4)* and *bla*_NDM-5_ genes on different ST. This strain of *E. coli* carries a large number of plasmids. The *tet(X4)*-positive *E. coli* carries different replicon types including Col, IncFIA, IncFIC, IncHI, IncI and IncX1, and harbouring multiple antimicrobial resistance genes, including *tet(X4)* and *bla*_NDM-5_ along with *aadA*, *bla*_TEM-1B_, *sul*, *floR* and *qnrS1 et, al.*.

## CONCLUSION

As far as we know, this study identified the ST5299 *E. coli* strain co-harboring *bla*_OXA-48_-like and *mcr-1*, and the ST5409 and ST877 *E. coli* strain co-harboring *bla*_NDM-5_ and *tet(X4)* gene cluster for the first time. Regardless of their low prevalence rate in environment-associated sources, the mobile plasmid-mediated resistance genes in such superbugs can pose a significant threat to public health. Therefore, continuous monitoring of such MDR bacteria in humans, animals, and the environment should be considered and to guide the deployment of public health interventions before clinical cases increase.

## MATERIALS AND METHODS

### Isolates collection

A total of 316 swine wastewater samples were collected from 29 pig farms across different regions in Shandong Province, China. At least one pig farm was sampled in each region, and 3 to 5 samples were taken from the manure collection pits of at least two pig houses per farm. Unfortunately, due to biosecurity control measures at the pig farms, sufficient samples could not be obtained in every region. The swine wastewater samples were incubated with LB broth without antibiotic selection. Then, the samples were cultured by inoculation onto MacConkey plates containing 2.0 mg/L meropenem and incubated for 18h at 37C. A red colony was selected from each plate for identification. Cultures were identified using MALDI-TOF MS Axima^TM^ (Shimadzu-Biotech Corp., Kyoto, Japan) and 16S rRNA sequencing. For the carbapenem-resistant isolates, five major carbapenemase resistance genes (*bla*_KPC_, *bla*_NDM_, *bla*_IMP_, *bla*_OXA-48_-like and *bla*_VIM_) were detected for in carbapenem-resistant isolates by PCR using previously described primers (32).

### Antimicrobial susceptibility testing

The MICs of 14 antibiotics (cefotaxime, ceftazidime, aztreonam, amikacin, gentamicin, ciprofloxacin, tetracycline, tigecycline, fosfomycin, sulfamethoxazole/trimethoprim, imipenem, meropenem, colistin, and florfenicol) for all recovery isolates were determined by agar dilution and interpreted according to the CLSI guidelines (CLSI, 2024). Susceptibility to colistin and tigecycline was tested via broth microdilution, and breakpoints for *Enterobacteriaceae* were interpreted following EUCAST (“European Committee on Antimicrobial Susceptibility Testing. Breakpoint tables for interpretation of MICs and zone diameters”) 2024 breakpoint criteria.. *E. coli* ATCC 25922 served as a quality control strain for susceptibility testing.

### Conjugation assay testing

To determine the transferability of the resistance genes, streptomycin resistant *E. coli* strain C600 was used as the recipient and the conjugation assay was performed using a filter mating method. Transconjugants were selected using MacConkey agar plates containing both 1.0 mg/L meropenem and 1,500 mg/L streptomycin. The identification of the transconjugants used PCR (33).

### WGS and phylogenetic analysis

Clonal relatedness of all of the *Enterobacteriaceae* was analyzed by ERIC-PCR as described previously (34). The genomic DNA of all Non-clonal *bla*_NDM_-positive *Enterobacteriaceae* was subjected to 250 bp paired-end WGS using the Illumina MiSeq system (35), and the paired-end Illumina reads were assembled by SPAdes v3.6.2.18 MLST (36), antibiotic resistance genes (ARGs), plasmid replicon types was analysed using the CGE server (https://cge.cbs.dtu.dk/services/).

The hosts and countries of 397 *E. coli* isolates were retrieved from NCBI (https://www.ncbi.nlm.nih.gov/pathogens) and the assembly genomes of the 397isolates downloaded from the NCBI database (as of Sep. 2024). All assembly genomes were used for core-genome alignments to produce a phylogenetic tree using the Parsnp software of the Harvest suite (37). In this pipeline, bases that have likely undergone recombination are removed using PhiPack and only columns passing a set of filters based on these criteria are considered reliable core-genome SNPs (38). The final set of core-genome SNPs was submitted to FastTree 2 for reconstructing a maximum likelihood phylogenetic tree using default parameters (39). The VCF file of all variants identified by Parsnp was then used to determine pair-wise single nucleotide variant distances between the core genomes of all strains. For the phylogenetic tree, a reference genome was randomly selected using the ‘-r!’ switch. The lineages of the phylogenetic tree were defined using rhierbaps version 6.0 (40). The heat map was generated using R 3.3.2 (R Foundation for Statistical Computing) and was used to construct the tree that was visualized using FigTree v1.4.2 and iTOL v4 (41). Generation of plasmid maps were performed with BRIG.

## DECLARATION OF COMPETING INTEREST

All authors approved the manuscript and gave their consent for submission and publication.

**Ying Chu:** Writing-original draft, Writing-review & editing. **Yujie Miao:** Data curation, Software. **Liya Huang:** Validation, Visualization. **Rong Wang:** Methodology, Visualization. **Yuxin Wang:** Data curation. **Fengting Liao:** Visualization. **Xiang Luo:** Validation. **Shuancheng Bai:** Conceptualization, Formal analysis, Project administration, Supervision.**Yubao Li:** Conceptualization, Resources

## ACKNOWLEDGMENTS

The author(s) declare that financial support was received for the research and/or publication of this article. This research was funded by the Development Program for Young Innovative Teams in Higher Education Institutions of Shandong Province (2024KJI019); The results of the Research Initiation Project for High-level Talents of Yulin Normal University in 2024 (G2024ZK13).

## DATA AVAILABILITY

All WGS data have been deposited in the NCBI ( Sequence Read Archive (SRA) submission: SUB15732648).

## Notes

### Competing Interest Statement

The authors have declared no competing interest.

### Summary of Updates

Updated the title, the preface, the results, and the reference citation format.

